# Patient-derived vascularized skin organoids unravel the role of systemic sclerosis fibroblasts in microvascular dysfunction

**DOI:** 10.64898/2025.12.03.691822

**Authors:** Amandine Pitaval, Lara Jobeili, Caroline Welsch, Stéphanie Combe, Anastasia Papoz, Laure Gibot, Matthieu Roustit, Jean-Luc Cracowski, Charles Coutton, Xavier Gidrol, Walid Rachidi

## Abstract

Systemic sclerosis (SSc) is a complex connective tissue disease characterized by excessive extracellular matrix deposition and endothelial dysfunction leading to skin fibrosis and vascular complications. To investigate the contribution of fibroblasts to SSc-associated vasculopathy, we generated 3D vascularized skin organoids incorporating patient-derived fibroblasts and healthy endothelial cells. Comparative analyses revealed that SSc fibroblasts alone induce microvascular alterations, including giant capillaries and early fibrotic responses at the secretome level. These findings demonstrate that fibroblasts are a central driver of vascular remodeling in SSc and highlight their key crosstalk with endothelial cells. This SSc-specific organoid model offers a valuable platform for studying disease mechanisms and therapeutic interventions without relying on animal models.

## Introduction

Systemic sclerosis (SSc) is a complex and connective tissue disease characterized by vascular alterations, inflammation and tissue fibrosis (Christopher P Denton 2016). Fibrosis in SSc arises from sustained fibroblast activation, resulting in excessive extracellular matrix deposition and the accumulation of alpha-smooth muscle actin (αSMA)-expressing myofibroblasts. These changes, which accumulate over time, result in tissue stiffening and progressive skin fibrosis (Wynn 2007). Importantly, the fibrotic process in SSc is preceded by early and profound microvascular dysfunction, a hallmark that is particularly prominent in the limited cutaneous form of the disease (Lc-SSc), This vasculopathy is characterized by endothelial dysfunction and distinctive structure abnormalities including capillary rarefaction, and dilation commonly referred to as “giant capillaries (Dinsdale et al. 2017). Beyond serving as an early hallmark, this microvascular dysfunction contributes directly to severe clinical manifestations of the disease, such as severe Raynaud’ phenomenon, digital ulcers, pulmonary hypertension, highlighting the central role of vascular pathology in SSc progression (Asano and Sato 2015),(Trojanowska 2010),(Chora et al. 2015).

The molecular mechanisms driving these clinical and pathological manifestations remain complex and incompletely understood. In vitro models of SSc have demonstrated that fibrosis can be induced by exogenous profibrotic signals such as TGFβ (Matei et al. 2019), by intrinsically activated SSc fibroblasts (Huang et al. 2020), or indirectly by disease-derived endothelial cells (Serratı et al. 2013) or keratinocytes (Russo et al. 2021) thought the activation of fibroblast. While these studies clearly establish fibroblasts as the central effectors of fibrosis, their role in microvascular remodeling remains poorly defined. In particular, direct cellular drivers of vascular dysfunction have not yet been clearly identified, although available evidence suggests that vascular alterations may be induced indirectly through exogeneous TGFβ-driven fibroblast activation (Kramer et al. 2022).

To address this gap, we developed reproducible vascularized human skin organoids comprising a dermal compartment containing primary human dermal fibroblasts, endothelial cells (HUVECs) and adipose-derived stem cells (ADSCs), together with an epidermis formed by primary human keratinocytes. Using this platform, we generated vascularized Lc-SSc skin organoids by replacing healthy fibroblasts with fibroblasts derived from Lc-SSc patients. By combining patient-derived fibroblasts with healthy endothelial cells in a 3D context, we tested whether Lc-SSc fibroblasts are sufficient to induce microvascular abnormalities, including the formation of giant capillaries and act as active drivers of pathological vascular remodeling in systemic sclerosis

## Results

### Vascularized full-thickness skin organoids display a dermis rich in ECM, a mature endothelial network and a differentiated epidermis

We generated and characterized vascularized skin organoids with N1-HDF (human primary fibroblasts derived from a healthy donor), GFP-expressing HUVECs, ADSCs and keratinocytes following the protocol described in the supplementary methods section (Figure 1A). For simplicity, only the origin of fibroblasts is indicated on the skin organoid images.

**Figure 1:**
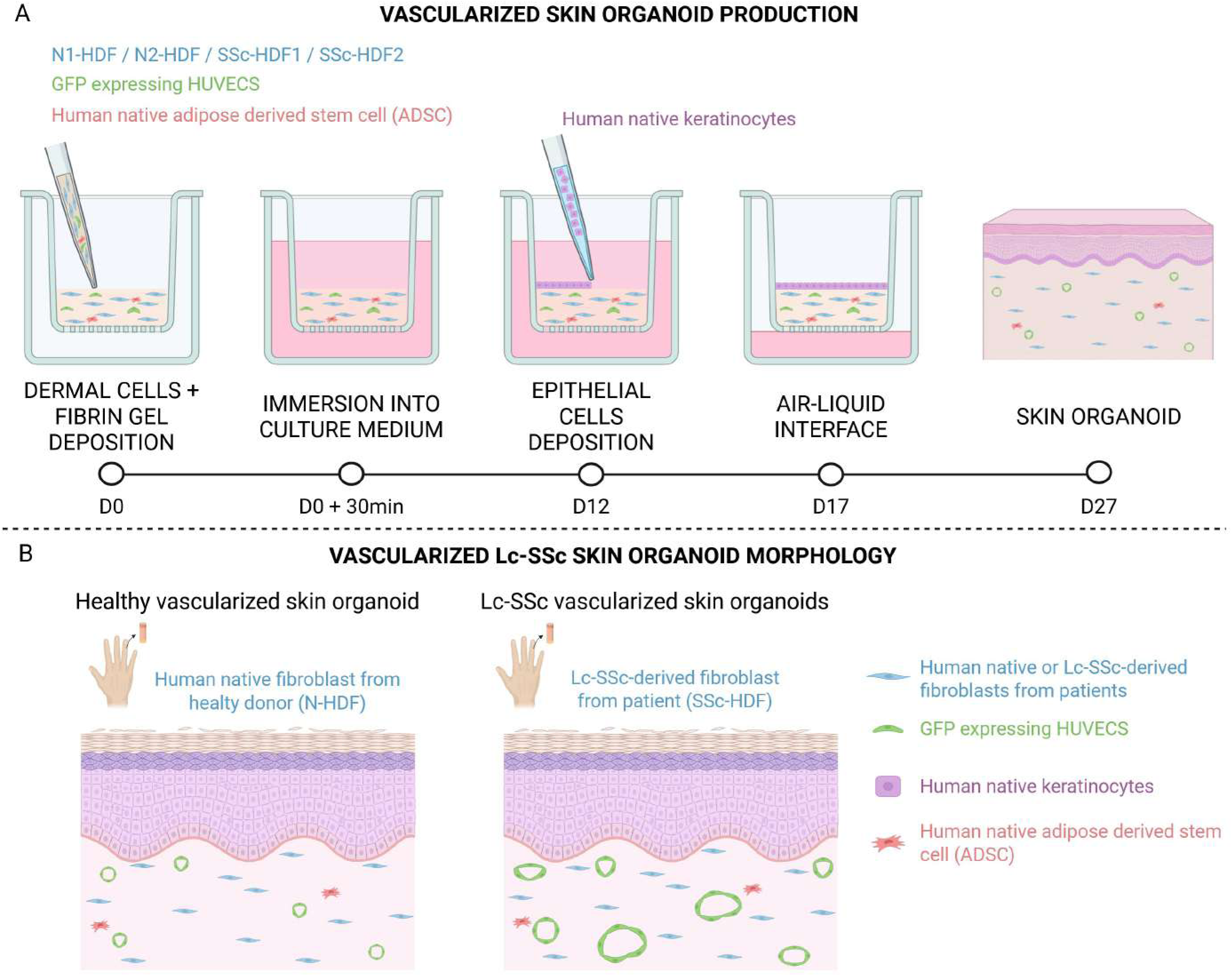
Graphical abstract illustrating the workflow for vascularized skin organoid production (A) and summarizing the pathological phenotype of Lc-SSc organoids with healthy donor skin organoids (B) (Created with BioRender.com. BioRender.com/f74t381)

After 27 days of culture, hematoxylin-eosin saffron (HES) staining of paraffin sections highlighted the morphology of vascularized skin organoids. The results showed that these organoids properly mimicked human native skin architecture (Figure 2A). Two distinct cellular compartments were distinguishable: (1) the dermis with low cell density, dense ECM and the presence of a developed endothelial network and (2) the epidermis that appeared dense, highly organized, pluri-stratified, and differentiated, with all cell layers (basal, spinous, and granular layers as well as the stratum corneum) similar to native human skin. Epidermal structure was assessed by analyzing filaggrin and cytokeratin 14 (CK14) expression. Filaggrin was appropriately localized to the upper layers of the epidermis consistent with terminal differentiation, with a distribution pattern comparable between native skin and skin organoids. CK14 expression was restricted to the basal layer in normal human skin, whereas a more diffuse pattern extending into suprabasal layers was observed in skin organoid (Figure 2B). The thickness of the epidermis in those organoids was 87 µm with a 95% CI of [81; 93] (Figure 2A/B), consistent with values reported for mammary skin (76.9 µm ±26.2 µm) (Oltulu et al. 2018). These results indicate proper basal cell proliferation and cell differentiation in the epidermis.

**Figure 2:**
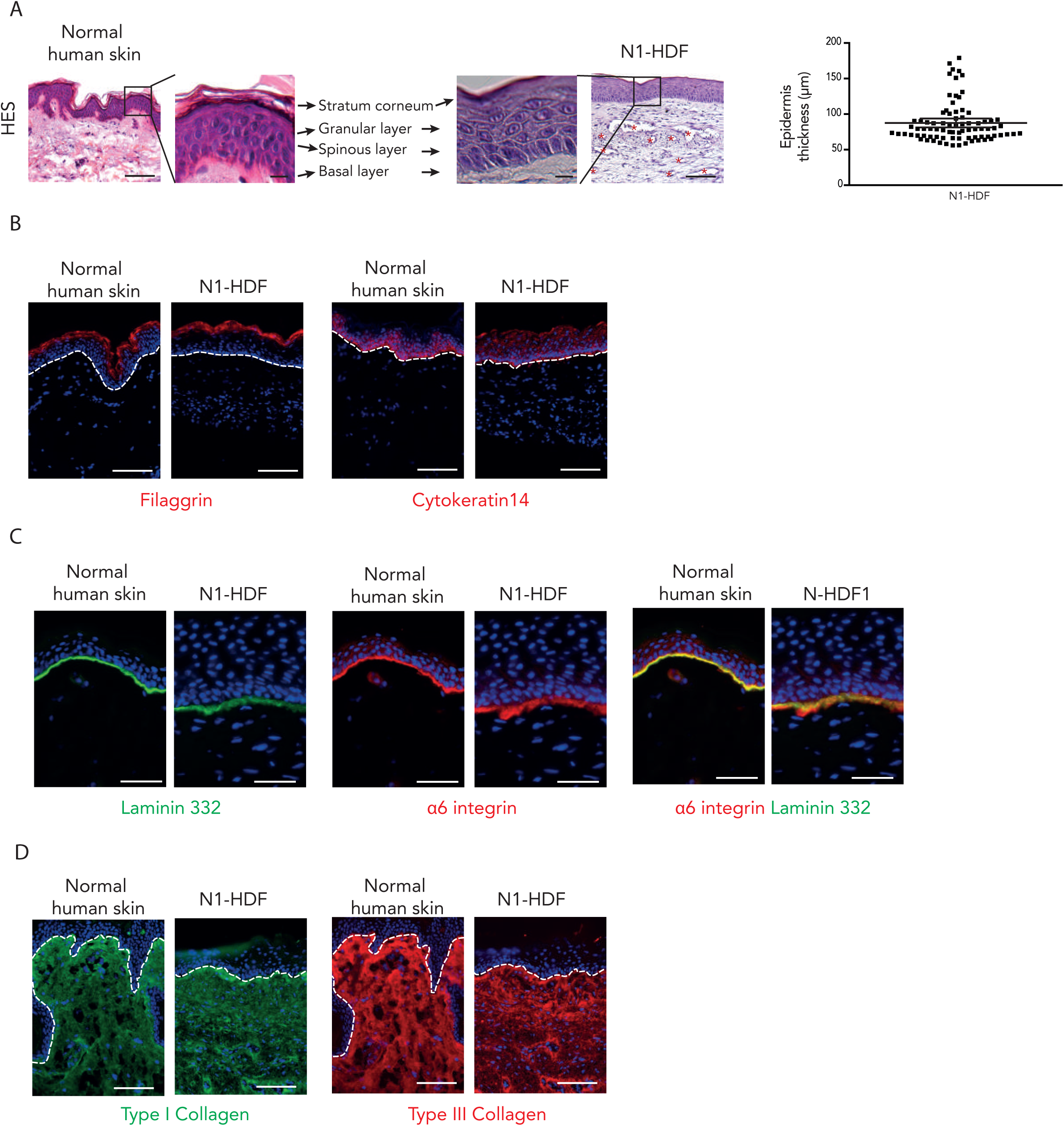
Characterization of complex physiological full-thickness vascularized skin organoids generated using healthy donor-derived fibroblast (N1-HDF), GFP-HUVECs, ADSCs and keratinocytes. To simplify image labeling, only the fibroblast origin (N1-HDF) is indicated. A/ Morphological features of normal human skin and skin organoids following hematoxylin-eosin-saffron (HES) staining of paraffin-embedded sections. Vascularized skin organoids were cultured for 27 days. HES staining of cross-sections showed a dermis containing endothelial structures (red stars) and a well-developed epidermis. Higher magnification of the epidermis showed complete stratification with the four expected cell layers: basal, spinous, granulous layers and stratum corneum. Scale bar: 25 µm. Right: quantification of epidermal thickness expressed in μm, measured from the basal layer to the granular layer. 80 independent fields from three independent experiments (two technical replicates per experiment) were analyzed. Scale bars: 100 µm. Graph values are presented as mean ± 95% CI. B/ Immunofluorescence staining of epidermal differentiation (Filaggrin: red) and basal keratinocyte proliferation (Cytokeratin 14: red). The dotted line indicates the dermal-epidermal junction (DEJ). Scale bars: 100 µm. C/ Immunofluorescence staining of DEJ proteins: laminin 332 (green) and α6 integrin (red). Scale bars: 50 µm. D/ Immunofluorescence staining of type I (green) and type III collagens (red) in the dermis. The dotted line indicates the DEJ. Scale bars: 100 µm. Nuclei were counterstained with DAPI (B, C, D).

Immunostaining of laminin 332 and α6 integrin, two markers of the dermal-epithelial junction (DEJ), confirmed the presence of an intact basement membrane. Their expression levels appeared qualitatively equivalent to that of native skin (Figure 2C).

Regarding the dermis composition, we focused on type I and type III collagen, two main components of the ECM secreted by fibroblasts (Figure 2D). Patterns of expression of type I and type III collagen were homogeneously distributed throughout the dermis and resembled the distribution observed in the native skin.

The generation and maturation of endothelial networks in vascularized skin organoids were both monitored with GFP-HUVECs using an inverted microscope. By D21 (during air-liquid interface), a dense, stable and interconnected endothelial network was detected (Figure 3A). The fluorescent labelling of CD31 (PECAM-1), a marker of endothelial cells, underlined the auto-organization of an endothelial network distributed in the whole dermis (Figure 3B). These capillaries contained a lumen (also observed in HES Figure 2A). The mean area of the endothelial lumen was 238 µm² with a 95% CI of [189; 288], representing 0.5% of the total dermis area (Figure 3C). The large dispersion was explained by the fact that vessels were cut either orthogonally or vertically into cross sections.

**Figure 3:**
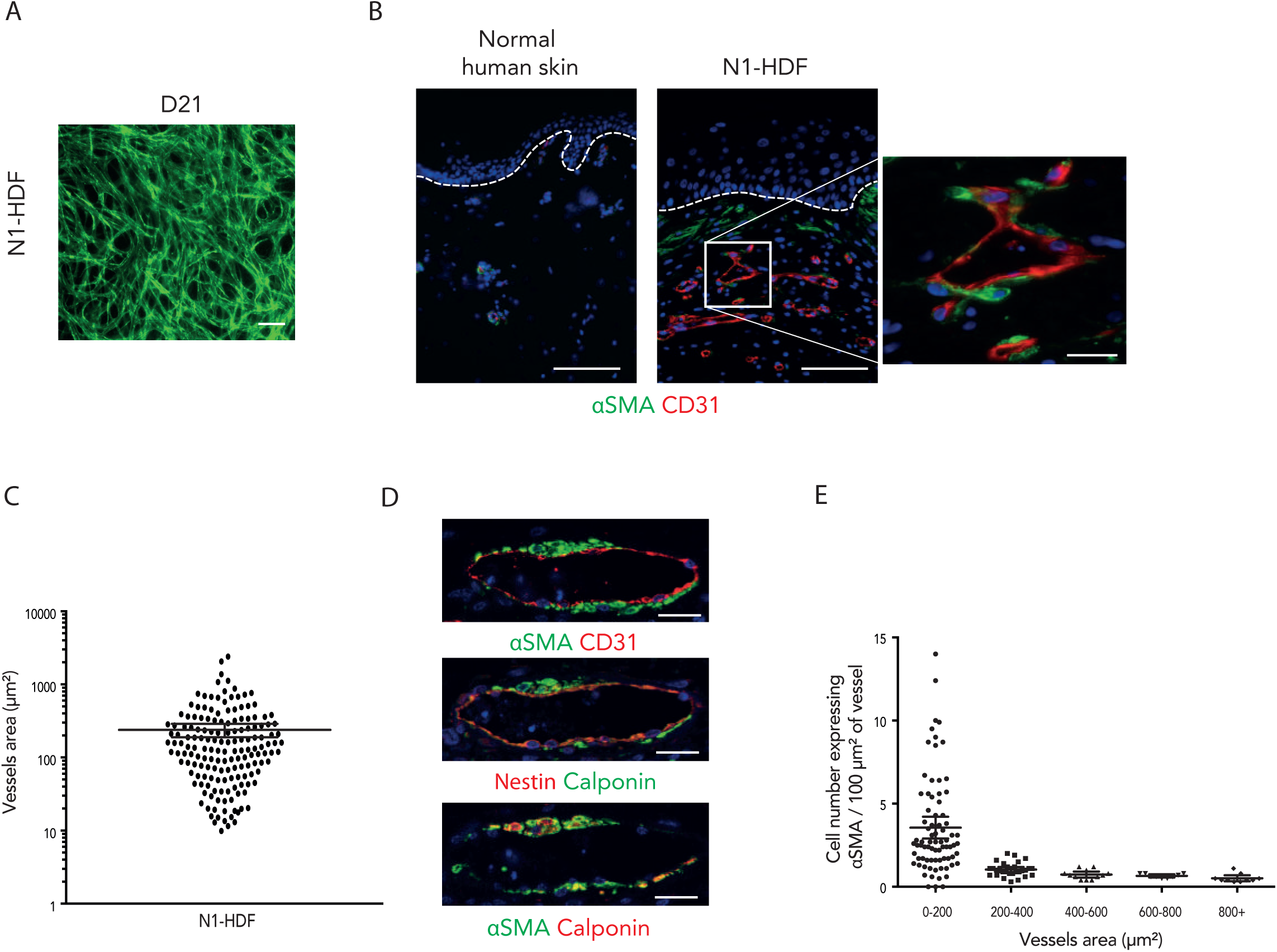
Characterization of the endothelial network in vascularized skin organoids. A/ Imaging of endothelial network development using GFP-HUVEC at day 21 (including seven days at the air-liquid interface) during skin organoid maturation. Bottom view of the skin organoid captured using an inverted microscope. Scale bar: 200 µm. B/ Immunofluorescence of vessel-like structures in vascularized skin organoids using CD31 labeling (red) and αSMA staining (green). Nuclei were counterstained with DAPI. Scale bars: 100 µm. The dotted line indicates the DEJ. The image on the right shows a higher magnification view of a vessel-like structure (area: 670 µm²). Scale bar: 25 µm. C/ Quantification of vessel lumen area (µm²) with a log10-transformed y-axis. Ten independent fields from three independent experiments (two technical replicates per experiment) were analyzed. Values are presented as mean ± 95% CI. D/ Three serial sections with three different stainings are shown from left to right: αSMA (green)/CD31 (red); αSMA (green)/Calponin (red); Calponin (green)/Nestin (red). Scale bars: 25 µm. E/ Distribution of pericyte-like cells as a function of vessel area.

Alpha smooth muscle actin (αSMA) is a marker of both myofibroblasts and pericytes. In the dermis, αSMA-positive cells were mainly located in the upper layer of the dermis attesting to the presence of myofibroblasts in the papillary dermis (Figure 3B). Moreover, some αSMA-positive cells were aligned along capillaries and seemed to be in physical contact with them, suggesting that they were pericyte-like cells (Figure 3B). To ensure that αSMA constitutes a suitable pericyte marker, a supplemental staining of Calponin (pericyte marker) was used on successive cross-sections (Figure 3D). The colocalization between αSMA and Calponin confirmed that the cells surrounding capillaries were pericytes-like cells.

Around each capillary, there was on average 2.3 αSMA-positive cell / 100 µm^2^ of vessel (data not shown). Interestingly, when the vessel area was sub-grouped by 200 µm², the vessel coverage of pericytes was 3.5 times higher on vessels whose area was below 200 µm² compared to larger vessels between 200 µm² and 400 µm² and more (Figure 3E).

Altogether, these observations suggest that vascularized skin organoids appear to reproduce several major structural characteristics of native human skin including epidermal organization, a preserved DEJ, a dermal compartment with homogeneous ECM content, myofibroblasts and matured vessel-like structures with lumens and surrounded pericytes.

### Vascularized skin organoids generated with patient-derived fibroblasts recapitulate some features observed in limited cutaneous systemic sclerosis skin

Healthy and limited cutaneous systemic sclerosis vascularized skin organoids were generated with GFP-HUVECs, primary human ADSCs, primary human keratinocytes, and either primary dermal human fibroblasts derived from a healthy donor (N2-HDF) or two Lc-SSC patients (SSc-HDF1 and SSc-HDF2) matched in sex and age (Table 2). Thus, in this part, we assess the impact of Lc-SSc fibroblasts from two Lc-SSC patients on vascular architecture.

**Table 1:**
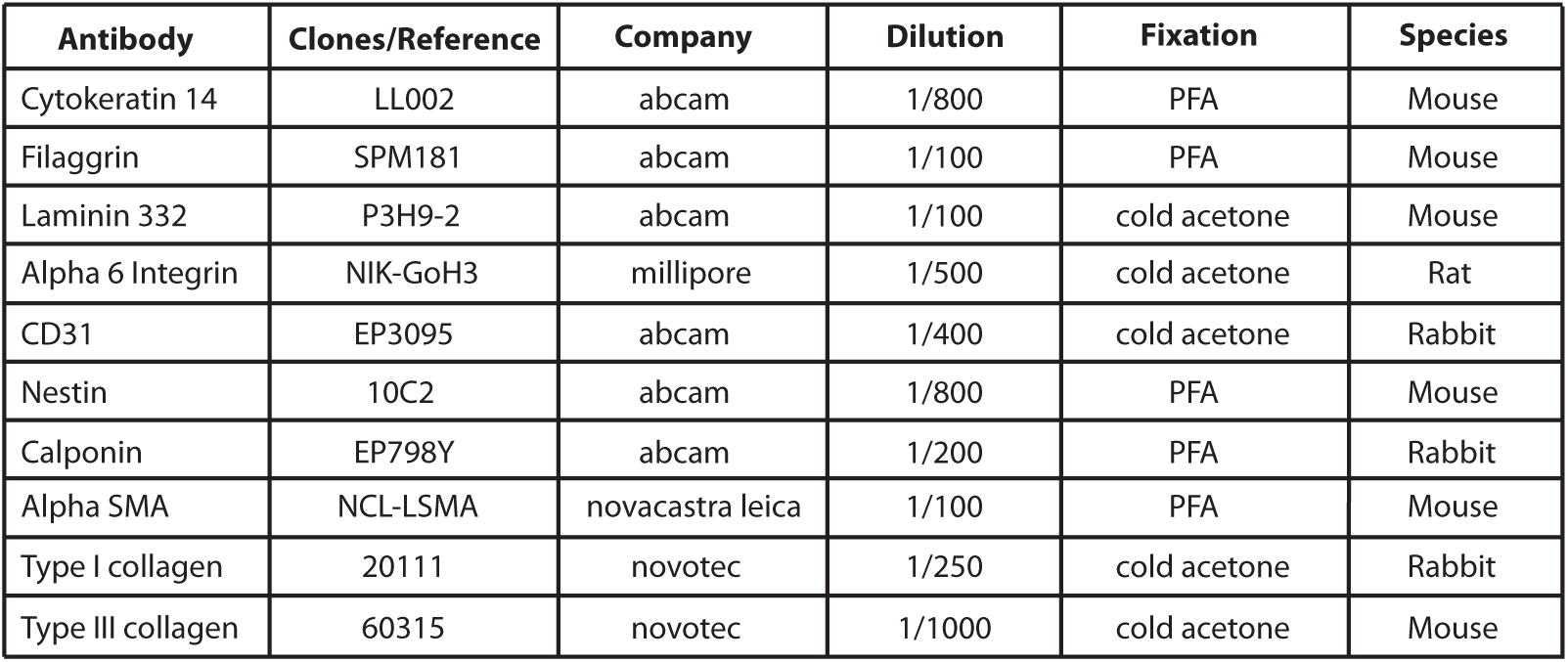
List of primary antibodies used in this study.

**Table 2:** Clinical features of the healthy donor and the two Lc-SSc (limited cutaneous systemic sclerosis) patients.

|  | N2-HDF | SSc1-HDF | SSc2-HDF |
| --- | --- | --- | --- |
| Age | 54 | 54 | 54 |
| Raynaud condition score (0-10) | 0 | 5 | 5 |
| Disease duration (years) | 0 | 14 | 23 |
| Modified Rodnan score (0-66)<br>(proximal interphalangeal joint) | 0 | 2 | 4 |
| Cutulo pattern | 0 | Early (few<br>megacapillaries) | Active (numerous<br>megacapillaries) |
| Videocapillaroscopy | Normal | Abnormal | Abnormal |

After 27 days of culture, HES staining of paraffin sections highlighted the particular morphology of Lc-SSc vascularized skin organoids (Figure 4). For both healthy and pathological conditions, two cellular compartments were distinguishable: (1) the dermis with low cell density, dense ECM and the presence of a rich endothelial network; and (2) the epidermis that appeared dense, highly organized, pluri-stratified, and well differentiated. The thickness of the epidermis of pathological skin organoids was measured on HES staining (Figure 4A). The quantification revealed a slight yet significant decrease in the epidermis thickness for SSc-HDF2 compared to healthy N2-HDF. To further analyze the homeostasis of the epidermis, we monitored terminal differentiation and proliferation using filaggrin and CK14 expression, respectively. We did not observe any quantitative or qualitative differences among all three conditions (Figure 4B). These data indicate that using fibroblasts from Lc-SSc patients did not affect skin architecture except for the epidermis thickness in SSc-HDF2.

**Figure 4:**
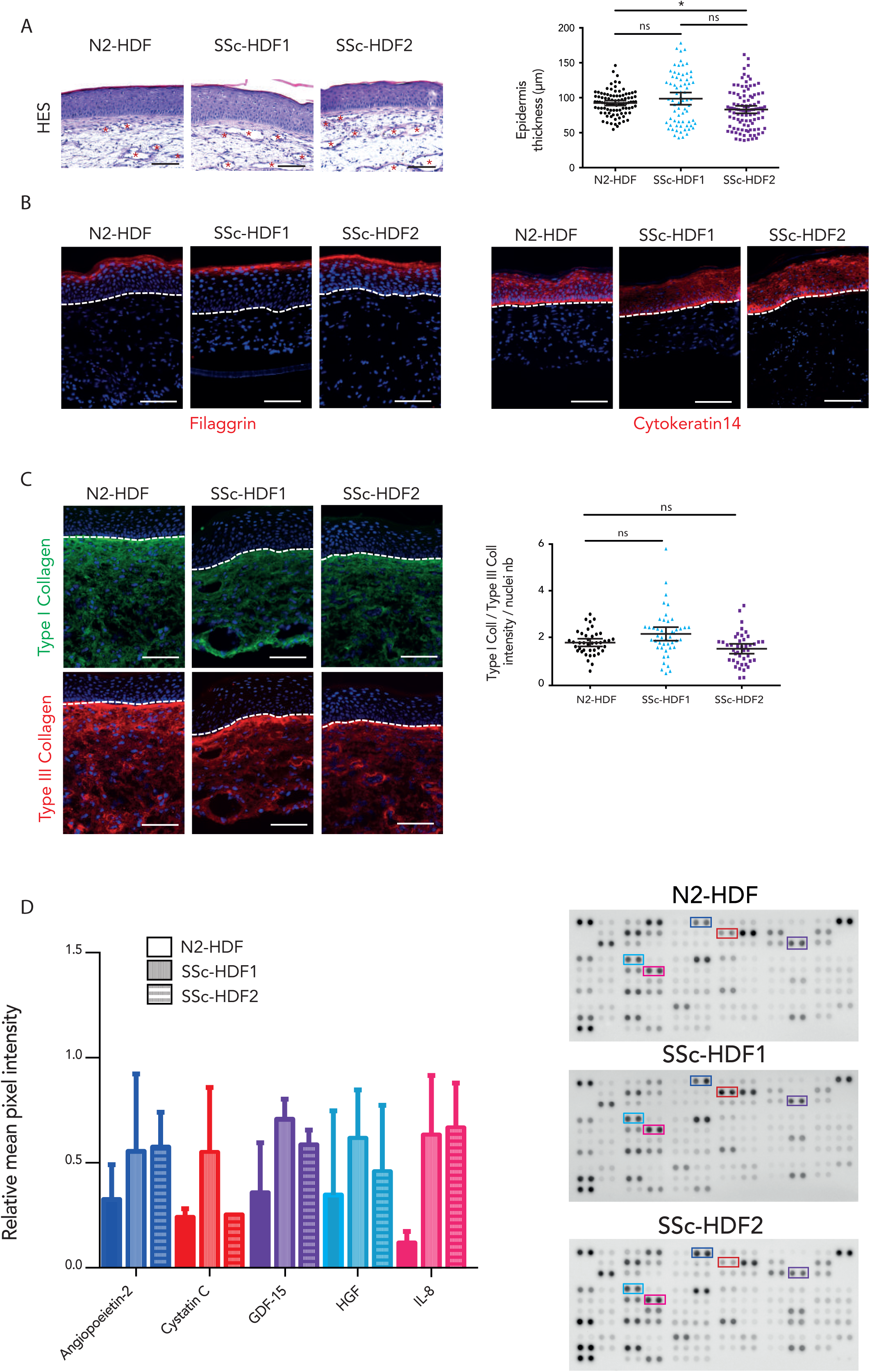
Characterization of complex pathological vascularized skin organoids generated using fibroblasts derived from two patients suffering from limited cutaneous systemic sclerosis (Lc-SSC). All skin organoids were generated with a combination of fibroblasts, GFP- HUVECs, adipose-derived stem cells (ADSCs) and keratinocytes. To simplify image labeling, only the fibroblasts origin is indicated (N2-HDF: healthy donor, SSc-HDF1: patient 1 with Lc-SSC, SSC-HDF2: patient 2 with Lc-SSC). HUVECs, ADSCs, and keratinocytes were not patient-derived from individuals suffering from Lc-SSC. A/ Morphological features after HES staining on paraffin-embedded sections of vascularized skin organoids generated with fibroblasts from one healthy donor (N2-HDF) and two patients with Lc-SSC (SSc-HDF1 and SSc-HDF2). Skin organoids were cultured for 27 days. Quantification of epidermal thickness was expressed in μm and measured from the basal layer to the granular layer. Ninety independent fields from three independent experiments (two technical replicates per experiment) were analyzed. Scale bars: 100 µm. Red stars indicate endothelial structures. B/ Immunofluorescence staining of epidermal differentiation (Filaggrin: red) and basal keratinocyte marker (Cytokeratin 14: red). The dotted line indicates the DEJ. Scale bars: 100 µm. C/ Immunofluorescence staining of type I (green) and type III collagens (red) in the dermis. Fluorescence intensity was quantified as an expression ratio normalized to dermal cell number. Fifteen independent fields from three independent experiments were analyzed. The dotted line indicates the DEJ. Scale bars: 100 µm. Nuclei were counterstained with DAPI (B, C, D). Graph values are presented as mean ± 95% CI (A, C). D/ Assessment of cytokine levels in the culture supernatants of vascularized skin organoids generated with fibroblasts derived from patients suffering from Lc-SSC. Graph representing a selection of the most differentially secreted cytokines between the healthy donor and the two Lc-SSc conditions. One membrane from each of the two biological replicates per condition is shown (mean ± SD). *p<0.05, **p<0.01, *** p<0.001, ****p<0.0001.

The accumulation of excess extracellular matrix components such as collagen defines fibrosis, one of the SSc features. We therefore compared the ratio of collagen type III/I (a marker of connective tissue pathology) in the pathological skin organoids. At the time of analysis (27 days of culture), it did not significantly change between pathological and normal conditions (Figure 4C). In addition, myofibroblasts expressing αSMA were mainly located in the upper layer of the dermis (Figure 5B). No significant difference was observed among all three conditions (data not shown).

**Figure 5:**
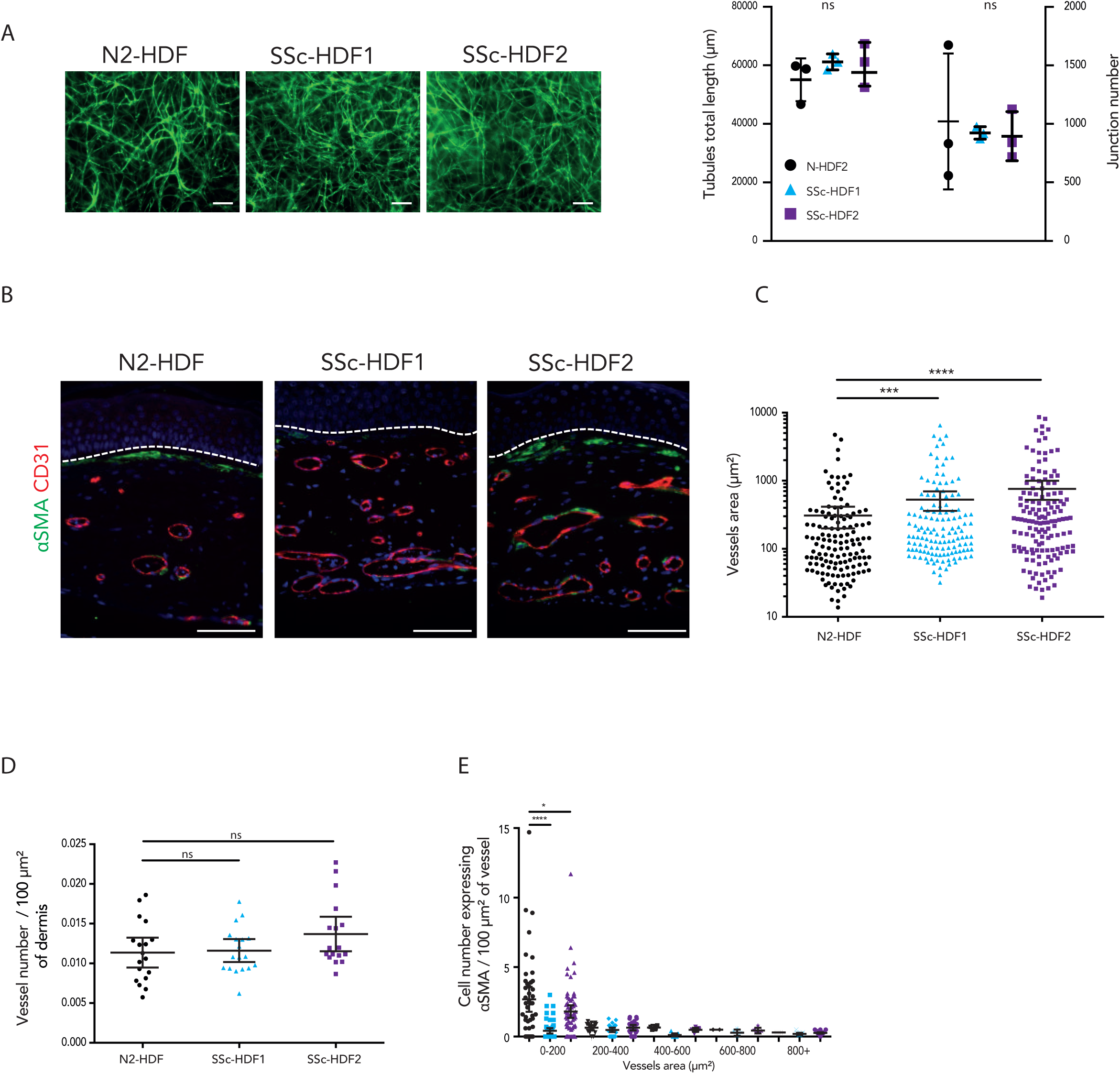
Characterization of the endothelial network in pathological vascularized skin organoids generated with SSc fibroblasts. A/ Endothelial network at day 21 (including 7 days at the air-liquid interface) during skin organoid maturation using GFP-HUVECs. Bottom view of the skin organoid captured using an inverted microscope. Scale bar: 200 µm. Quantification of total tubule length and junction number was performed at this time point (three independent experiments). B/ Immunofluorescence staining of vessel-like structures in vascularized skin organoids using CD31 (red) and αSMA (green) staining. Nuclei were counterstained with DAPI. Scale bars: 100 µm. Lower panels show higher magnification view of vessel- like structures. Scale bars: 50 µm. The dotted line indicates the DEJ. C/ Quantification of vessel lumen area (µm, log10-transformed y-axis) for healthy donor (N2-HDF) and the two Lc-SSC patients (SSc-HDF1 and SSc-HDF2). Ten independent fields from three independent experiments (two technical replicates per experiment). Values are presented as mean ± 95% CI. Graph on the right: Distribution of pericyte-like cell as a function of vessel area. D/ Graph representing the vessel number normalized to the dermis area. Results were expressed as number of vessels per 100 µm2 of dermal area E/ Distribution of pericyte-like cell as a function of vessel area. Quantification was performed on ten independent fields from three independent experiments (two technical replicates per experiment). Values are presented as mean ± 95% CI. *p < 0.05, **p < 0.01, *** p < 0.001, ****p < 0.0001.

In Lc-SSc disease, several cytokines and chemokines have been implicated in the induction of fibrosis (Shima 2021). Their dysregulation plays a central role in initiating and maintaining inflammatory responses resulting in angiogenesis alterations and fibrosis (Randone et al. 2017). Using cytokine array, we assessed the relative level of human cytokine and chemokine secretions in the case of pathological skin organoids compared to the healthy control. Figure 4D shows a selection of the most secreted and differentially secreted cytokines between the healthy donor and the two pathological Lc-SSc. In the majority of cases, an increase of cytokine secretion was observed in both pathological conditions compared to N2-HDF. All cytokines were represented in the form of a heat map in Figure S1 and in a table (Table 3).

**Table 3:**
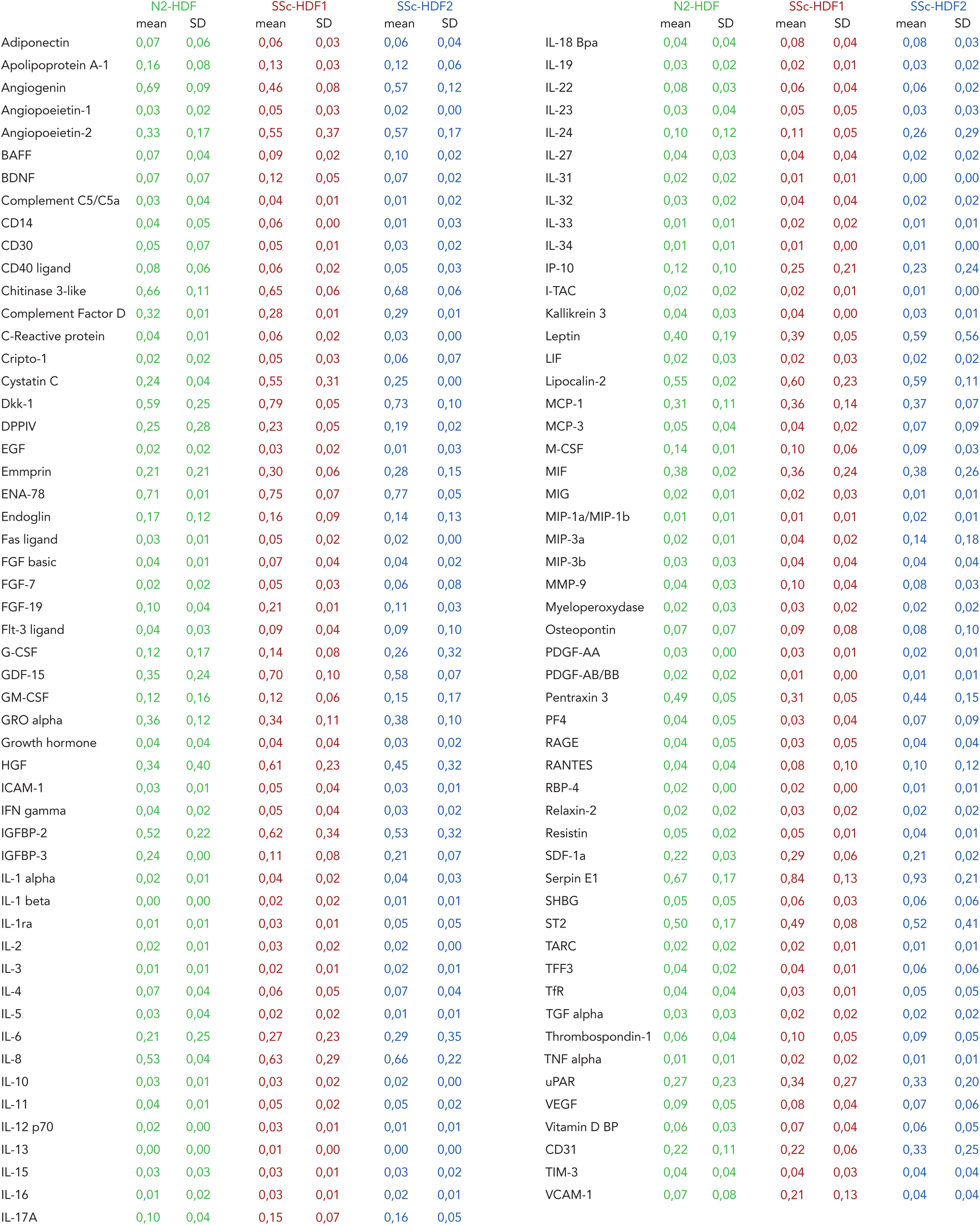
Cytokine levels in skin organoids generated with fibroblasts derived from Lc-SSc patients. Data represent mean ± SD of all the secreted cytokines for the healthy donor and the two pathological Lc-SSc organoids.

|  | N2-HDF |  | SSc-HDF1 |  | SSc-HDF2 |  |  | N2-HDF |  | SSc-HDF1 |  | SSc-HDF2 |  |
| --- | --- | --- | --- | --- | --- | --- | --- | --- | --- | --- | --- | --- | --- |
|  | mean | SD | mean | SD | mean | SD |  | mean | SD | mean | SD | mean | SD |
| Adiponectin | 0,07 | 0,06 | 0,06 | 0,03 | 0,06 | 0,04 | IL-18 Bpa | 0,04 | 0,04 | 0,08 | 0,04 | 0,08 | 0,03 |
| Apolipoprotein A-1 | 0,16 | 0,08 | 0,13 | 0,03 | 0,12 | 0,06 | IL-19 | 0,03 | 0,02 | 0,02 | 0,01 | 0,03 | 0,02 |
| Angiogenin | 0,69 | 0,09 | 0,46 | 0,08 | 0,57 | 0,12 | IL-22 | 0,08 | 0,03 | 0,06 | 0,04 | 0,06 | 0,02 |
| Angiopoeitin-1 | 0,03 | 0,02 | 0,05 | 0,03 | 0,02 | 0,00 | IL-23 | 0,03 | 0,04 | 0,05 | 0,05 | 0,03 | 0,03 |
| Angiopoeitin-2 | 0,33 | 0,17 | 0,55 | 0,37 | 0,57 | 0,17 | IL-24 | 0,10 | 0,12 | 0,11 | 0,05 | 0,26 | 0,29 |
| BAFF | 0,07 | 0,04 | 0,09 | 0,02 | 0,10 | 0,02 | IL-27 | 0,04 | 0,03 | 0,04 | 0,04 | 0,02 | 0,02 |
| BDNF | 0,07 | 0,07 | 0,12 | 0,05 | 0,07 | 0,02 | IL-31 | 0,02 | 0,02 | 0,01 | 0,01 | 0,00 | 0,00 |
| Complement C5/C5a | 0,03 | 0,04 | 0,04 | 0,01 | 0,01 | 0,02 | IL-32 | 0,03 | 0,02 | 0,04 | 0,04 | 0,02 | 0,02 |
| CD14 | 0,04 | 0,05 | 0,06 | 0,00 | 0,01 | 0,03 | IL-33 | 0,01 | 0,01 | 0,02 | 0,02 | 0,01 | 0,01 |
| CD30 | 0,05 | 0,07 | 0,05 | 0,01 | 0,03 | 0,02 | IL-34 | 0,01 | 0,01 | 0,01 | 0,00 | 0,01 | 0,00 |
| CD40 ligand | 0,08 | 0,06 | 0,06 | 0,02 | 0,05 | 0,03 | IP-10 | 0,12 | 0,10 | 0,25 | 0,21 | 0,23 | 0,24 |
| Chitinase 3-like | 0,66 | 0,11 | 0,65 | 0,06 | 0,68 | 0,06 | I-TAC | 0,02 | 0,02 | 0,02 | 0,01 | 0,01 | 0,00 |
| Complement Factor D | 0,32 | 0,01 | 0,28 | 0,01 | 0,29 | 0,01 | Kallikrein 3 | 0,04 | 0,03 | 0,04 | 0,00 | 0,03 | 0,01 |
| C-Reactive protein | 0,04 | 0,01 | 0,06 | 0,02 | 0,03 | 0,00 | Leptin | 0,40 | 0,19 | 0,39 | 0,05 | 0,59 | 0,56 |
| Cripto-1 | 0,02 | 0,02 | 0,05 | 0,03 | 0,06 | 0,07 | LIF | 0,02 | 0,03 | 0,02 | 0,03 | 0,02 | 0,02 |
| Cystatin C | 0,24 | 0,04 | 0,55 | 0,31 | 0,25 | 0,00 | Lipocalin-2 | 0,55 | 0,02 | 0,60 | 0,23 | 0,59 | 0,11 |
| Dkk-1 | 0,59 | 0,25 | 0,79 | 0,05 | 0,73 | 0,10 | MCP-1 | 0,31 | 0,11 | 0,36 | 0,14 | 0,37 | 0,07 |
| DPPIV | 0,25 | 0,28 | 0,23 | 0,05 | 0,19 | 0,02 | MCP-3 | 0,05 | 0,04 | 0,04 | 0,02 | 0,07 | 0,09 |
| EGF | 0,02 | 0,02 | 0,03 | 0,02 | 0,01 | 0,03 | M-CSF | 0,14 | 0,01 | 0,10 | 0,06 | 0,09 | 0,03 |
| Emmprin | 0,21 | 0,21 | 0,30 | 0,06 | 0,28 | 0,15 | MIF | 0,38 | 0,02 | 0,36 | 0,24 | 0,38 | 0,26 |
| ENA-78 | 0,71 | 0,01 | 0,75 | 0,07 | 0,77 | 0,05 | MIG | 0,02 | 0,01 | 0,02 | 0,03 | 0,01 | 0,01 |
| Endoglin | 0,17 | 0,12 | 0,16 | 0,09 | 0,14 | 0,13 | MIP-1a/MIP-1b | 0,01 | 0,01 | 0,01 | 0,01 | 0,02 | 0,01 |
| Fas ligand | 0,03 | 0,01 | 0,05 | 0,02 | 0,02 | 0,00 | MIP-3a | 0,02 | 0,01 | 0,04 | 0,02 | 0,14 | 0,18 |
| FGF basic | 0,04 | 0,01 | 0,07 | 0,04 | 0,04 | 0,02 | MIP-3b | 0,03 | 0,03 | 0,04 | 0,04 | 0,04 | 0,04 |
| FGF-7 | 0,02 | 0,02 | 0,05 | 0,03 | 0,06 | 0,08 | MMP-9 | 0,04 | 0,03 | 0,10 | 0,04 | 0,08 | 0,03 |
| FGF-19 | 0,10 | 0,04 | 0,21 | 0,01 | 0,11 | 0,03 | Myeloperoxidase | 0,02 | 0,03 | 0,03 | 0,02 | 0,02 | 0,02 |
| Flt-3 ligand | 0,04 | 0,03 | 0,09 | 0,04 | 0,09 | 0,10 | Osteopontin | 0,07 | 0,07 | 0,09 | 0,08 | 0,08 | 0,10 |
| G-CSF | 0,12 | 0,17 | 0,14 | 0,08 | 0,26 | 0,32 | PDGF-AA | 0,03 | 0,00 | 0,03 | 0,01 | 0,02 | 0,01 |
| GDF-15 | 0,35 | 0,24 | 0,70 | 0,10 | 0,58 | 0,07 | PDGF-AB/BB | 0,02 | 0,02 | 0,01 | 0,00 | 0,01 | 0,01 |
| GM-CSF | 0,12 | 0,16 | 0,12 | 0,06 | 0,15 | 0,17 | Pentraxin 3 | 0,49 | 0,05 | 0,31 | 0,05 | 0,44 | 0,15 |
| GRO alpha | 0,36 | 0,12 | 0,34 | 0,11 | 0,38 | 0,10 | PF4 | 0,04 | 0,05 | 0,03 | 0,04 | 0,07 | 0,09 |
| Growth hormone | 0,04 | 0,04 | 0,04 | 0,04 | 0,03 | 0,02 | RAGE | 0,04 | 0,05 | 0,03 | 0,05 | 0,04 | 0,04 |
| HGF | 0,34 | 0,40 | 0,61 | 0,23 | 0,45 | 0,32 | RANTES | 0,04 | 0,04 | 0,08 | 0,10 | 0,10 | 0,12 |
| ICAM-1 | 0,03 | 0,01 | 0,05 | 0,04 | 0,03 | 0,01 | RBP-4 | 0,02 | 0,00 | 0,02 | 0,00 | 0,01 | 0,01 |
| IFN gamma | 0,04 | 0,02 | 0,05 | 0,04 | 0,03 | 0,02 | Relaxin-2 | 0,02 | 0,02 | 0,03 | 0,02 | 0,02 | 0,02 |
| IGFBP-2 | 0,52 | 0,22 | 0,62 | 0,34 | 0,53 | 0,32 | Resistin | 0,05 | 0,02 | 0,05 | 0,01 | 0,04 | 0,01 |
| IGFBP-3 | 0,24 | 0,00 | 0,11 | 0,08 | 0,21 | 0,07 | SDF-1a | 0,22 | 0,03 | 0,29 | 0,06 | 0,21 | 0,02 |
| IL-1 alpha | 0,02 | 0,01 | 0,04 | 0,02 | 0,04 | 0,03 | Serpin E1 | 0,67 | 0,17 | 0,84 | 0,13 | 0,93 | 0,21 |
| IL-1 beta | 0,00 | 0,00 | 0,02 | 0,02 | 0,01 | 0,01 | SHBG | 0,05 | 0,05 | 0,06 | 0,03 | 0,06 | 0,06 |
| IL-1ra | 0,01 | 0,01 | 0,03 | 0,01 | 0,05 | 0,05 | ST2 | 0,50 | 0,17 | 0,49 | 0,08 | 0,52 | 0,41 |
| IL-2 | 0,02 | 0,01 | 0,03 | 0,02 | 0,02 | 0,00 | TARC | 0,02 | 0,02 | 0,02 | 0,01 | 0,01 | 0,01 |
| IL-3 | 0,01 | 0,01 | 0,02 | 0,01 | 0,02 | 0,01 | TFF3 | 0,04 | 0,02 | 0,04 | 0,01 | 0,06 | 0,06 |
| IL-4 | 0,07 | 0,04 | 0,06 | 0,05 | 0,07 | 0,04 | TfR | 0,04 | 0,04 | 0,03 | 0,01 | 0,05 | 0,05 |
| IL-5 | 0,03 | 0,04 | 0,02 | 0,02 | 0,01 | 0,01 | TGF alpha | 0,03 | 0,03 | 0,02 | 0,02 | 0,02 | 0,02 |
| IL-6 | 0,21 | 0,25 | 0,27 | 0,23 | 0,29 | 0,35 | Thrombospondin-1 | 0,06 | 0,04 | 0,10 | 0,05 | 0,09 | 0,05 |
| IL-8 | 0,53 | 0,04 | 0,63 | 0,29 | 0,66 | 0,22 | TNF alpha | 0,01 | 0,01 | 0,02 | 0,02 | 0,01 | 0,01 |
| IL-10 | 0,03 | 0,01 | 0,03 | 0,02 | 0,02 | 0,00 | uPAR | 0,27 | 0,23 | 0,34 | 0,27 | 0,33 | 0,20 |
| IL-11 | 0,04 | 0,01 | 0,05 | 0,02 | 0,05 | 0,02 | VEGF | 0,09 | 0,05 | 0,08 | 0,04 | 0,07 | 0,06 |
| IL-12 p70 | 0,02 | 0,00 | 0,03 | 0,01 | 0,01 | 0,01 | Vitamin D BP | 0,06 | 0,03 | 0,07 | 0,04 | 0,06 | 0,05 |
| IL-13 | 0,00 | 0,00 | 0,01 | 0,00 | 0,00 | 0,00 | CD31 | 0,22 | 0,11 | 0,22 | 0,06 | 0,33 | 0,25 |
| IL-15 | 0,03 | 0,03 | 0,03 | 0,01 | 0,03 | 0,02 | TIM-3 | 0,04 | 0,04 | 0,04 | 0,03 | 0,04 | 0,04 |
| IL-16 | 0,01 | 0,02 | 0,03 | 0,01 | 0,02 | 0,01 | VCAM-1 | 0,07 | 0,08 | 0,21 | 0,13 | 0,04 | 0,04 |
| IL-17A | 0,10 | 0,04 | 0,15 | 0,07 | 0,16 | 0,05 |  |  |  |  |  |  |  |

Altogether, we did not detect any changes in collagen deposition in the pathological models compared to the healthy control. Nevertheless, cytokines and chemokines associated with Lc-SSc fibrosis were markedly increased, suggesting that their secretion precedes the fibrotic process. Some of these cytokines may also impact the vascular network, reflecting aspects of vasculopathy. Based on these observations, we next examined endothelial network development in all conditions.

### Vascularized limited cutaneous systemic sclerosis skin organoids contain giant capillaries

During culture at the air liquid interface until day 21, endothelial network was stable and interconnected (Figure 5A). Although it appeared denser with pathological fibroblasts, measurements of total tubule length and junction number showed no significant difference (Figure 5A).

HES staining of paraffin sections and immunofluorescence using CD31 highlighted the presence of a developed endothelial network and the presence of larger vessels in pathological conditions (Figure 4A, Figure 5B). Strikingly, the quantification on CD31-labelled sections showed that although the number of vascular structures did not change significantly between healthy and pathological skin organoids, the lumen area was significantly increased with Lc-SSc fibroblasts (SSc-HDF1: 528 µm² with a 95% CI of [360; 695]; SSc-HDF2:759 µm² with a 95% CI of [522; 997]) compared to healthy donor fibroblasts (N2-HDF: 307 µm² with a 95% CI of [199; 416]) (Figure 5C, D). These larger vessel-like structures are classically called “giant capillaries”. In both conditions, they could reach a lumen surface of more than 6500 µm² (Figure 5C).

Around each vascular structure, the number of pericytes observed along vessels was quantified for each condition (Figure 5E). When the vessel area was classified into subgroups, we noticed that on vessels smaller than 200 µm^2^, the pericyte coverage was significantly reduced in the presence of Lc-SSC fibroblasts. There were 2.7 αSMA-positive cells / 100 µm² of vessel for N2-HDF, 1.2 for SSc-HDF1, and 1.8 for SSc-HDF2. Overall, larger vessels with an area of more than 200 µm^2^ were surrounded by very few pericytes, whatever the origin of the fibroblasts.

Altogether, we show for the first time that Lc-SSc fibroblasts substantially impact the organization of the endothelial network, leading to the formation of giant capillaries observed in the dermis of Lc-SSc patients. Furthermore, these results suggest that these patient-derived vascularized skin organoids, as they recapitulate some critical features of the disease, could be used as a relevant model of systemic sclerosis to better understand the etiology of the disease regarding the vasculopathy or to screen drugs that could help patients.

## Discussion

We generated vascularized skin organoids by co-culturing fibroblasts, endothelial cells, keratinocytes, and ADSCs in a fibrin hydrogel. ADSCs, which can differentiate into pericyte-like cells, stabilized the endothelial networks and contributed to extracellular matrix production supporting both vascular organization and overall skin architecture (Song et al. 2016),(Andrée et al. 2019),(Rohringer et al. 2014),(Frueh et al. 2017) (Metral et al. 2014). Under these conditions, keratinocytes self-organized into a stratified epidermis reminiscent of native human skin, while endothelial cells formed lumenized vessels surrounded by pericyte-like cells, recapitulating key features of the microvasculature.

We next aimed at generating vascularized skin organoids that recapitulate key features of limited cutaneous systemic sclerosis (Lc-SSc). The disease follows a well-defined sequence of clinical manifestations. It begins with endothelial cell injury and microvascular dysfunction, including abnormal capillary morphology (dilated or giant capillaries) and reduced capillary density (Dinsdale et al. 2017). These vascular alterations precede B- and T-cell activation with exacerbated chemokine and cytokine production. These immune events lead to tissue inflammation and hypoxia and fibroblast proliferation. Late clinical features include collagen accumulation, fibrosis, progressive thickening and hardening, and in severe cases, ulceration (Ko et al. 2023).

Incorporation of Lc-SSc fibroblasts in our study recapitulated microvascular alterations characterized by abnormal vessel morphology, including giant capillaries, a hallmark of early vascular dysfunction. Vascular defects have also been reported following exogenous TGFβ exposure, which impairs angiogenesis and leads to microvascular rarefaction, a later-stage process, highlighting the impact of TGFβ signaling in vascular integrity (Kramer et al. 2022). In our organoid, we observed an earlier vascular event, likely due to the origin of the fibroblasts, which were derived from the lateral digits of patients, a region with pronounced vascular abnormalities but minimal fibrosis. These observations suggest that these Lc-SSC fibroblasts were sufficient to induce microvascular defects, including the formation of giant capillaries, closely resembling patient capillaroscopy patterns (Table 2).

Interestingly, we also showed that pericytes were preferentially associated with smaller vessels, consistent with their role in stabilizing capillaries and regulating vessel diameter capillary blood flow (Siekmann 2023). Moreover, pericyte coverage was reduced on small vessels in pathological conditions, suggesting altered endothelial/pericyte interactions in this disease.

The mechanisms driving microvascular alterations in Lc-SSc remain incompletely understood. They could include endothelial apoptosis, impaired endothelial proliferation, altered endothelial cell migration or endothelial-to-mesenchymal transition (EndoMT), all of which contribute to abnormal vessel morphology, as observed in SSc (Chora et al. 2015) (Manetti et al. 2017) (Romano, Rosa, and Fioretto 2024). Additional factors such as pericyte detachment, vascular basement membrane composition and integrity, and dysregulated angiogenic signaling, may further exacerbate vessel dilatation and structural abnormalities, providing a mechanism framework for the vascular alteration seen in Lc-SSC (Siekmann 2023) (Renaud et al. 2024).

In our model, we did not observe features of late-stage fibrotic features such as increased collagen accumulation, indicating that it predominantly captures early pathogenic events rather than established fibrosis. This absence of overt matrix deposition is consistent with the early vascular alterations observed in our organoids. Nevertheless, we detected increased secretion of cytokines and soluble factors known to contribute to inflammation, vasculopathy and early fibrotic signaling such as GDF15, IL-8, HGF, and Angiopoietin-2 (Randone et al. 2017) (Kawaguchi et al. 2002) (Carvalheiro et al. 2020). These findings suggest that soluble factors may promote early vascular dysfunction and inflammatory activation prior overt fibrosis remodeling.

The lack of fibrosis in our model contrasts with a 3D skin model reporting fibrotic remodeling induced by SSc fibroblasts at later stages (Huang et al. 2020). In that study, collagen deposition and myofibroblast activation were enhanced by exogeneous TGFβ supplementation, likely reflecting a more advanced profibrotic phase of SSc. In contrast, our model appears to recapitulate an earlier stage in which fibroblasts-driven vascular dysfunction occurs without triggering overt fibrosis.

Moreover, in our study, the epidermis remained also largely unaffected by the presence of Lc-SSc fibroblasts, likely because keratinocytes were derived from a young and healthy donor. Bryon *et al* showed that dermal fibroblasts from SSc patients can communicate with keratinocytes via exosomes to trigger a type I interferon response in the epidermis without causing detectable morphological changes in keratinocytes, indicating a dermis to epidermis crosstalk (Bryon et al. 2025).

Conversely, keratinocytes from SSc patients display functional abnormalities in 3D culture models, including altered differentiation, reduced proliferation and activation, which are also observed in fibrotic skin or scars (Russo et al. 2021) (N. Aden et al. 2008) (Nima Aden et al. 2010) (Nikitorowicz-Buniak et al. 2014). These patient-derived keratinocytes can additionally influence the dermis, including differential expression of extracellular matrix, highlighting a dermis - epidermis crosstalk (McCoy et al. 2017),(Nima Aden et al. 2010). Notably there is currently no evidence that SSc keratinocytes directly contribute to microvascular alterations. Including Lc-SSc keratinocytes in our model would therefore be valuable to investigate whether they could affect vascularization.

Expanding the number of Lc-SSc donors in future studies will be essential to determine whether the observed vascular phenotypes are conserved across SSc subsets and to better capture inter-individual variability, a hallmark of the disease (Buechler et al. 2021).

## Conclusion

Here, we present a complex 3D vascularized human skin organoid model generated with fibroblasts derived from Lc-SSc patients, who recapitulate some earlier features of the disease, such as secretion of cytokines preceding fibrosis, and microvasculopathy (Figure 1B). This bioengineered vascularized skin using primary fibroblasts derived from patients, offer a robust model to unravel the role of Lc-SSc patient-derived fibroblasts in endothelial dysfunction. Finally, as it recapitulates early manifestations of the disease, this model could be very useful to screen drugs that could reduce or delay later deleterious symptoms of scleroderma disease.

## Materials and Methods

### Tissue harvesting and primary cell culture

Human primary fibroblasts called N1-HDF and human primary keratinocytes were extracted from healthy young subjects. Skin tissue explants were obtained from surgically discarded female breast specimens. N1-HDF used for development and optimization of skin organoids were isolated as previously described (Auxenfans et al. 2012) and cultured in Dulbecco’s modified Eagle medium (DMEM) supplemented with 10% FBS (Pan Biotech). After separation of the epidermis from the dermis, keratinocytes were isolated as previously described and grown in KSFM culture medium (Auxenfans et al. 2012).

Primary human pathological fibroblasts were isolated from skin biopsies harvested from diseased skin on the lateral face of the finger from two females with limited cutaneous systemic sclerosis (SSc-HDF1 and SSc-HDF2) and one age- and sex-matched healthy control subject (N2-HDF). Human primary fibroblasts called N2-HDF came from a healthy, age- and gender-matched donor. Fibroblasts were isolated using the explant method. The biopsies used for the culture of fibroblasts were immediately placed in Cytomax culture medium (Adgenix®). The sample was placed in a single-use petri dish, divided into several pieces using a scalpel, cut into small fragments, and placed in a CO_2_ oven. The cells were monitored daily with an inverted microscope. When the growth appeared sufficient, the cell pellets were recovered in PBS with a cryoprotector (DMSO) for freezing in nitrogen. All research protocols about the collection and use of human cells derived from skin biopsies were approved by the French ministry (DC-2008- 735, DC-2008-444). Pathological samples and N2-HDF were collected during the TRP clinical trial (NTC03211325). All subjects gave written informed consent, and studies were performed in accordance with ethical guidelines.

ADSCs (adipose-derived stem cells) were extracted from adipose tissue discarded from abdominoplasty of anonymous healthy patients (biopredic ref GRA001) as previously described (Bunnell et al. 2008).

The endothelial cells were GFP-expressing HUVECs (human umbilical vein endothelial cells) (cAP-0001GFP Angioproteomie) cultivated according to manufacturer recommendations.

Each cell type was cultured in a CO_2_ incubator at 37 °C, and the appropriate culture medium was changed three times per week before confluence was reached. For passage, cells were rinsed with PBS solution twice and incubated with trypsin-EDTA 0.05% (Thermo Fisher Scientific, Waltham, MA, USA) for 5 min. They were subsequently incubated with a 10% FBS solution and centrifuged at 300 g for 5 min prior to resuspension in fresh culture media and subsequent culture.

### In vitro, 3D vascularized human skin organoid preparation

Cell-laden hydrogel preparation and deposition:

For all experiments, a 2.5% (w/v) fibrinogen (F8630 Sigma–Aldrich) solution containing aprotinin (A6279 Sigma–Aldrich) and 2.5 mM CaCl_2_ (C8106 Sigma–Aldrich) was prepared.

To establish vascularized dermal equivalents, experiments were carried out as follows (Figure 1A):

Fibroblasts (0.9 x 10^6^ cells), GFP-HUVECs (0.4 x 10^6^ cells) and ADSCs (0.3 x 10^6^ cells) were added to 1 ml of fibrinogen solution.

After the addition of thrombin (T4648 Sigma–Aldrich) at a concentration of 1 U/ml, 360 µl of cell-laden hydrogel were immediately and carefully pipetted and deposited into each culture chamber (ref 3460 Transwell Corning) located in a 12-well plate and placed at 37 °C in an incubator containing 5% CO_2_ for 30 min. Two ml of a mixture of 1:1 medium composed of DMEM (ref 31966047 DMEM Gibco, Life Technologies Ldt. Paisley, UK) supplemented with 10% FBS, 100 units/ml penicillin, and 100 µg/ml streptomycin and CnT-ENDO (Cellntec, Berne, Switzerland) were then subsequently applied to the bottom of each insert and 500 µl to the top. During the incubation period (12 days), 80 µg/ml ascorbic acid (A8960-5g Sigma–Aldrich, St Louis, USA) and 10 ng/ml EGF (EGF; R&D Systems, Minneapolis, MN, USA) were added to the cell culture medium to allow the formation of a dermal equivalent.

On day 12, human primary keratinocytes at passage 1 were harvested, and 45 000 keratinocytes were seeded onto the dermal equivalent. Each skin organoid was cultured in immersed conditions for 5 days in a green-adapted medium(Black et al. 2005), changing the medium three times a week. On day 17, the skin organoid was raised at the air-liquid interface. To do so, the insert was removed from the 12-well plate and placed into a 12-well insert holder (657110 Dominique Dutscher, Brumath, France). Four milliliters of a simplified medium composed of DMEM supplemented with 8 mg/ml of bovine serum albumin (A2153-50G Sigma–Aldrich), 80 µg/mL L-ascorbic acid, and antibiotics was added below the insert so that the bottom of the organoid was in contact with the medium. The insert was cultured for 10 more days, changing the media three times a week to trigger the formation of a full-thickness skin equivalent.

Samples were harvested and either fixed in 4% neutral-buffered formalin overnight and embedded in paraffin after dehydration and xylene incubation or directly embedded in Tissue Tek compound and frozen at -20 °C for histological and immuno-histological analysis. Three replicates were produced under biological conditions. Native human skin was also added for histological analysis as a reference.

### Live endothelial network

The endothelial network from GFP-expressing HUVECs developement was monitored at different time points during skin organoid formation, and images were captured using an AxioObserver optical microscope (ZEISS Axio Observer Z1). Image processing and analysis were performed using ImageJ software with the MBF plugin for microscopy (http://www.macbiophotonics.ca/imagej/, Research Service Branch, US National Institute of Health, United States), and the vascular network was quantified automatically by Angioquant software(Niemisto et al. 2005).

### Histology and immunohistological analyses

To assess the histological morphology of the skin organoids, paraffin-embedded formalin-fixed samples or frozen OCT samples were cut into 5 μm and 8 μm sections, respectively. After dewaxing and rehydration, sections were stained with hematoxylin, eosin, and saffron (HES staining) on paraffin-embedded sections. For immunofluorescence, labelling was performed either on paraffin-embedded or frozen OCT sections. Air-dried 8 µm cryo-sections were either incubated in 4% PFA solution for 15 min then in 0,1% Triton X-100 (Sigma–Aldrich) for 4 min or incubated in cold acetone solution for 10 min, followed by 3% BSA PBS incubation for 1 h to block nonspecific binding sites. Then, the slides were incubated with the primary antibodies listed in Table 1 diluted in 1,5% BSA in PBS overnight at 4 °C followed by Cy3- or Cy5-conjugated secondary antibodies (Jackson ImmunoResearch) for 30 min at RT. Nuclear counterstaining using DAPI was carried out according to a routing protocol. For αSMA / CD31 and Calponin / Nestin labelling, paraffin-embedded sections were treated with heat-mediated antigen retrieval treatment prior to incubation with 3% BSA PBS before the addition of primary and secondary antibodies. Image acquisition was performed using a Zeiss microscope (ZEISS Axio Imager Z1) for immunofluorescence experiments.

### Image analysis and processing

Image processing and analysis were performed using ImageJ. The living epidermal thickness, including all layers from the basal layer to stratum granulosum except the stratum corneum, was measured as the distance between the basement membrane to the stratum corneum on HES staining and expressed in µm on at least six fields per image and at least four images per sample. For analysis of type I and III collagen of the dermis, the fluorescently positive area, after setting a threshold, was automatically detected and segmented from other pixels and measured and normalized with the stromal nuclei number (except nuclei from vessel structure). Type I and III collagen intensities were analyzed in same field. The result was expressed as the ratio of type III collagen to type I collagen (type III/type I). Regarding CD31-positive vessel analysis, the area of each vessel was manually delineated and measured. The results represent the area of the vessel lumen and are expressed in µm^2^. The vessel area and the vessel number were normalized to the dermis area.

For αSMA expressing cells analysis, positive cells were counted around each CD31-positive vessel and subsequently normalized by the corresponding vessel area. The values were multiplied by 100 to get the number of pericytes per 100 µm^2^ of vessel. Finally, for DEJ (dermal epidermal junction), α6 integrin and laminin 332 were analyzed following the same process. The positive fluorescent area was automatically detected after setting a specific threshold for each labelling and the batch of images. The results were expressed as intensity relative to the length of the DEJ (in µm manually measured).

### Cytokine array

We used a human proteome profiler human XL cytokine array (ARY022B, R&D systems, Minneapolis, USA) according to the manufacturer’s instruction. Conditioned media from the four mature skin organoids (comprising either N1-HDF, N2-HDF, SSc-HDF1 or SSc-HDF2) were collected at the air-liquid interface stage. Each kit contained four membranes and each spotted in duplicate with 105 different cytokine-specific antibodies. Arrays were imaged using a Chemidoc^TM^ MP system (Bio-rad, Hercules, CA, USA). Pixel intensities of individual spots were quantified using Image J software (NIH, Bethesda, USA). For each antibody, the intensity of the two duplicated spots was first averaged. Background intensity was then subtracted and the resulting values were normalized to the mean intensity of the positive reference spots located at three corners of each membrane. For normalization, the mean intensity of the positive controls was set to 1, and all cytokine signals were expressed relative to this value. Two biological replicates were performed and signal intensities corresponding to the same antibody were finally averaged.

### Statistical analysis

To compare multiple groups with one grouping variable, one-way ANOVA with multiple comparisons and Tuckey’s correction was performed. Regarding the vessel area analysis, a non-parametric test was used because the data were not normally distributed. Statistical analyses were performed using GraphPad Prism 6 (GraphPad Software Inc., La Jolla, CA, USA). Data are presented as mean ± 95% confidence interval (CI). P values < 0.05 were considered statistically significant and are indicated throughout the manuscript as follows: *p < 0.05, **p < 0.01, ***p < 0.001 and ****p < 0.0001.

### Data and materials availability

Upon request, the data generated in this study are available from the corresponding authors.

## Supporting information

Relative cytokine levels in skin organoids generated with fibroblasts derived from Lc-SSc patients

## Conflict of Interest

The authors declare that they have no conflicts of interest

## Acknowledgments

This work was supported by ANR grant (ANR-18-CE17-0017), by the French National Research Agency under the Investissements d’Avenir program (ANR-15-IDEX-02), by “CEA DRF-impulsion”, and “CEA inflexion OoC” grants, and by ASF (Association Sclerodermique de France).

We gratefully acknowledge Delphine Freida and Frederique Kermarrec for ADSC cell extraction from adipose tissue. We also thank Rebecca Powell for her assistance with proofreading the manuscript.

---

**Supplementary Figure 1:** Relative cytokine levels in skin organoids generated with fibroblasts derived from Lc-SSc patients. Heat map representing all secreted cytokines for the healthy donor and the two pathological Lc-SSc organoids.

## Notes

### Competing Interest Statement

The authors have declared no competing interest.

### Summary of Updates

Figures reordered; some updates in the text

