## Supplementary figures and images for "Patient-derived vascularized skin organoids unravel the role of systemic sclerosis fibroblasts in microvascular dysfunction"

### Relative cytokine levels in skin organoids generated with fibroblasts derived from Lc-SSc patients

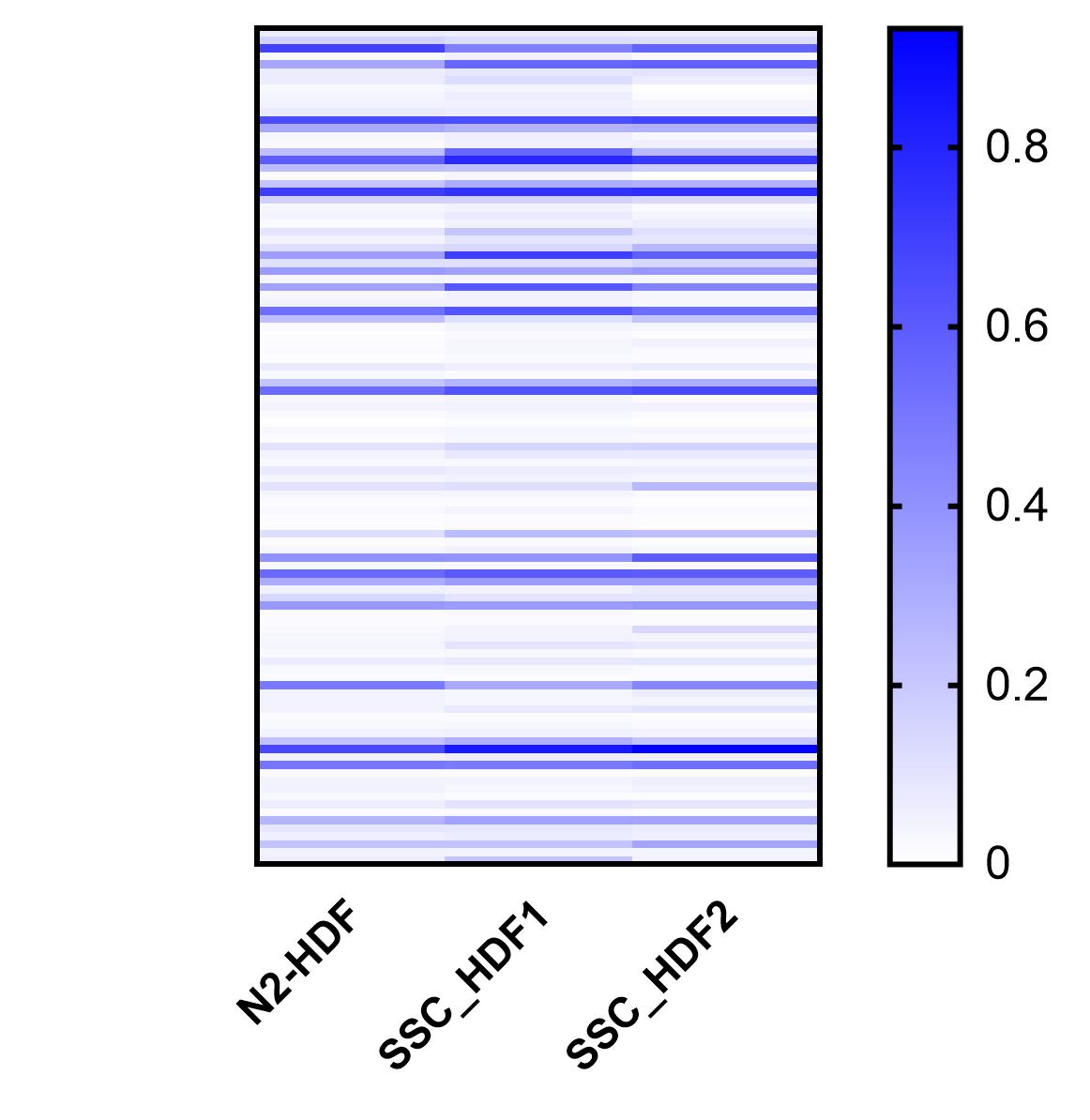
